# Genetic deletion and pharmacologic inhibition of FcER1 reduces renal injury in mouse models of diabetic nephropathy

**DOI:** 10.1101/2022.09.10.507433

**Authors:** Priyanka Rashmi, Andrea Alice Silva, Tara Sigdel, Izabella Damm, Ana Luisa Figueira Gouvêa, Suneil Koliwad, Vighnesh Walavalkar, Samy Hakroush, Minnie M. Sarwal

**Author notes:** Corresponding Author: Minnie M. Sarwal, MD, PhD, Medical Director, Kidney and Pancreas Transplant Program, Division of Multi-Organ Transplantation, Department of Surgery, University of California, San Francisco, 513 Parnassus Ave, Med Sciences Bldg, Room S1268, San Francisco, CA 94143. Authors contributed equally to the study (Priyanka Rashmi, Andrea Alice Silva).

## Abstract

FcER1 forms a high affinity multimeric cell-surface receptor for the Fc region of immunoglobulin E (IgE) and controls the activation of mast cells and basophils. Antigen binding and cross-linking of FcER1 associated IgE induces several downstream signaling pathways that result in diverse outcomes. Canonical signaling through IgE-FcER1 has been related to allergic responses, however, recent studies have identified that their function in mast cell and basophils contribute to other pathogenic conditions such as cancer and diabetes. Previous studies have demonstrated that FcER1 protein is upregulated in advanced diabetic kidney disease (DKD) making it a targetable molecule for the treatment of DKD. This study presents evidence that loss of FcER1 signaling reduces proteinuria and renal injury in two pre-clinical mouse models of diabetes. Mice deficient for *fcer1* are protected from streptozotocin mediated induction of proteinuria and display reduced fibrosis and mast cell infiltration in kidney. Furthermore, inhibition of FcER1 signaling with an antibody directed against the γ-subunit reduces proteinuria in a spontaneous model of type II diabetes. Our results show significant reduction of proteinuria and tissue damage in pre-clinical DKD models demonstrating the potential of FcER1 inhibitory approaches for developing new therapies in DKD.

## Introduction

Diabetic kidney disease (DKD) is a common co-morbidity associated with diabetes and the leading cause of end stage renal disease (ESRD) in developed countries (60.6% United States Renal Data System. 2021) [1]. DKD develops in approximately 30% of patients with type I and 40% patients with type II diabetes [2]. DKD It is a microvascular complication of diabetes, characterized by hyperfiltration leading to several structural alternations in multiple kidney compartments. The earliest consistent change is thickening of glomerular basement membrane paralleled by capillary and tubular basement membrane thickening. Other glomerular changes include mesangial matrix expansion, loss of endothelial fenestrations, loss of podocytes and foot process effacement. This results in global glomerulosclerosis and tubulointerstitial fibrosis. It is clinically defined as persistent albuminuria of an albumin-to-creatinine ratio above 30mg/g, with a progressive decline in kidney function [3]. Pathophysiology of DKD is driven by hyperglycemia and hypertension. Hyperglycemia is known to cause kidney hypertrophy by generating advanced glycation end products (AGEs), oxidative injury and hypoxia. These pathways lead to the activation of many growth factors such as insulin like growth factor 1 (IGF-1), transforming growth factor-β (TGFβ), epidermal growth factor (EGF) among others that mediate the progression of DKD. In addition, hyperglycemia can cause cellular injury that can trigger the release of proinflammatory factors including cytokines such as tissue necrosis factor-α (TNFα) and interleukin-1 (IL-1), adhesion molecules and damage-associated molecules (DAMPs). Infiltration of inflammatory cells that are activated in association with diabetes also contributes to structural changes in virtually all areas of kidney. Renal infiltration of macrophages and the resulting cytokine release creates a cycle with further monocyte and macrophage recruitment leading to inflammation associated structural alterations. Other cells such as mast cells also infiltrate the tubule-interstitium and release inflammatory factors. The rate of loss of estimated glomerular filtration (eGFR) has been demonstrated to be associated with the magnitude of macrophage infiltration and mast cell degranulation [4]. Currently there are no targeted therapies for DKD, and treatments are largely confined to diet and lifestyle changes and renin-angiotensin system blockades leaving large residual risk in DKD [5]. Over the last decade, omics technologies have been widely applied to study the systemic and local renal molecular changes to identify novel biomarkers and promising therapeutic targets in kidney disease [6]. A previous study performed microarray on PBMCs from a cohort of diabetic patients and non-diabetic controls to identify immunomodulatory genes that are significantly altered. These genes were mapped to transcriptionally altered genes in the glomeruli of patients with type II diabetes with biopsy proven diabetic nephropathy (publicly available data extracted from Gene Expression Omnibus (GEO). Gene expression of the gamma subunit of the high-affinity receptor of IgE, *Fcε receptor 1* (*fcer1*) in PBMCs was associated with the more advanced stages of diabetes and diabetic nephropathy as well as glomerular FcER1γ expression was higher in diabetic kidney [7]. FcER1 is constitutively expressed on mast cells and basophils [8]. IgE activates mast cells by binding to FcER1 and plays a critical role in mediating allergic responses [9]. Recent studies have also demonstrated a role for IgE in activating macrophages and T cells and collectively these responses contribute to inflammation associated with obesity and diabetes [10]. While many studies have been conducted to assess the role of IgE receptor and IgE targeting cells such as mast cells in the pathophysiology of diabetes and obesity, their role in the development and progression of kidney injury remains unclear. Genetic deficiency and pharmacological stabilization of mast cells has been shown to reduce diet-induced obesity and diabetes in mice [11]. Furthermore, deletion of *fcer1* has been described to improve glucose tolerance in mice with diet induced obesity [10] and anti FcER1 antibody administration delays the onset of type I diabetes in NOD mice [12]. However, diabetes associated with renal injury and function has not been explored in any of the above-mentioned models. In this study, we aimed to test the hypothesis that FcER1 mediates the progression of DKD, at least in part, and can be a therapeutic target in DKD for inhibiting or delaying disease progression. Therefore, we investigated the protective effects of FcER1 inhibition in two mouse models for type I (streptozotocin, STZ) and type II diabetes (leptin receptor mutatnt mice, db/db) to confirm the validity of FcER1 as a therapeutic target in DKD. STZ treatment in mice deficient for *fcer1* due to the gamma subunit deletion caused reduced proteinuria and histological renal injury to that in wild type mice. Furthermore, inhibition of FcER1 with an anti FcER1 antibody (MAR-1) in a spontaneous diabetes mouse model (db/db) delayed the progression of DKD.

## Materials and Methods

### Animals

All animals were housed in a controlled environment with a 12h light and 12h dark cycle with free access to water and a standard laboratory diet. All animal experiments were conducted under the approval of UCSF Institutional Animal Care and Usage Committee. The C57BL/6J wild type mice (*fcer1g*^*+/+*^) were purchased from Jackson Laboratory (Cat No. 000664, Bar Harbor, ME). The *fcer1γ*^-/-^ mice were purchased from Taconic Biosciences (Cat No. 583, USA). BKS.Cg-+Leprdb/+Leprdb/OlaHsd (db/db) mice were procured from the Jackson Lab (Cat No. 000642).

### Induction of hyperglycemia with Streptozotocin

Hyperglycemia was induced in wild type C57/BL6J and the *Fcer1g*^*-/-*^ mice according to the published protocol from NIDDK Diabetic Complications Consortium (DiaComp). Briefly, 10-week-old male mice were administered two doses of streptozotocin (STZ) (Sigma, 150 mg/kg body weight) in 50mM sodium citrate buffer, pH 4.5, intraperitoneally on two consecutive weeks (Figure 1A). Glucose levels from tail blood were measured with an Accu-Check glucometer (Roche Life Sciences). Animals presenting with blood glucose levels >350 mg/dL for two consecutive weeks were considered diabetic. 21 wild type and 23 *fcer1g*^*-/-*^ mice were used across four experiments and body weight, food and water intake, blood glucose, urine volume were recorded. At the end of each experiment serum and kidney tissue were collected as described.

**Figure 1.**
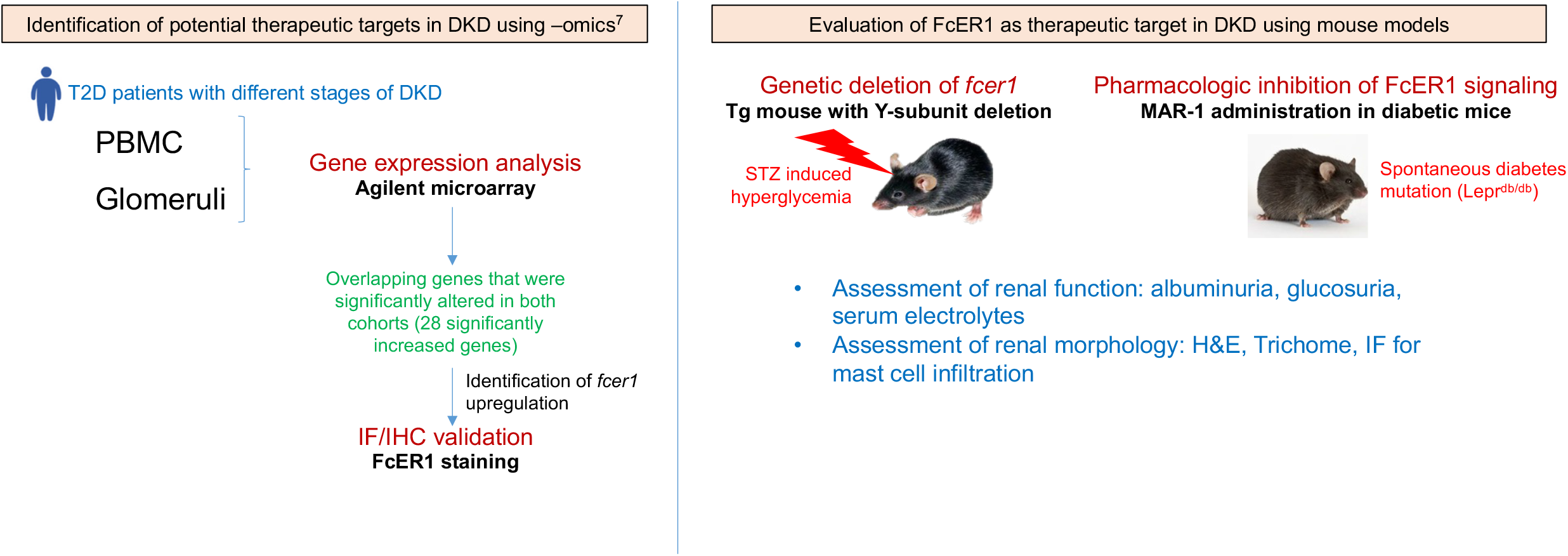
Study design. Increased expression of *fcer1* gene was identified in a -omics study in patients with advanced stage diabetic kidney disease (DKD) and showed elevation of the mRNA transcript and protein in renal tissue. This study uses two different mouse models of hyperglycemia (induced with streptozotocin mediated beta cell destruction and spontaneous development of diabetes in a mouse model with mutations in leptin receptor gene) and evaluates the disruption of FcER1 receptor mediated signaling as a therapeutic approach for DKD.

### Inhibition of FcER1 signaling with MAR-1 administration in db/db mice

10–16-week-old *<u>BKS.Cg-+Leprdb/+Leprdb/OlaHsd (db/db)</u>* mice were intraperitonially injected (IP) weekly with FcER1 alpha monoclonal antibody MAR-1 (Cat No. 16-5898-85, Ebioscience) or IgG isotype control (Cat No. 16-4888-85, Invitrogen) diluted with sterile saline (50ug, 50mg/kg) for six weeks. These experiments were performed at Melior Biosciences and UCSF Sarwal lab in collaboration with Dr. Suneil Koliwad. At the beginning and the end of study, blood glucose levels were measured, and 24-hour urine samples were collected using metabolic chambers. Bodyweights of animals were recorded on weekly basis.

### Blood and Urine collection

Mice were anesthetized with isoflurane. Blood was obtained from the retro-orbital plexus and centrifuged for 10 min at 3000rpm at RT. Aliquots of serum were made and stored at -80ºC until further use. For urine collection, the mice were acclimated to single housing metabolic cages (Techniplast, Italy) for 3 days. A powder diet of the same composition as regular laboratory diet was freely provided. Urine was collected for 4 hours and 24 hours for analysis. Urine was spun at 3000 rpm for 10 minutes to remove any food contamination and supernatant was collected. Aliquots of urine were made and stored at -80° until further analysis.

### Biochemical measurements

At the beginning and the end of study, the blood glucose levels were measured with glucometers and levels reported as mg/dL. Glucometers were calibrated prior to each measurement. Blood (<5 µL) was acquired from a tail snip and directly applied to a glucose test strip. Urine albumin levels was quantified using mouse albumin ELISA kit (cat. #E99-134, Bethyl Laboratories, Texas, USA). Urine creatinine was measured by creatinine kit (#MAK080, Sigma-Aldrich, MO, USA). Both the assays used the same samples and were performed on the same day. The urine albumin excretion was expressed as the albumin/creatinine ratio (uACR) in ug/mg. Urinary albumin and creatinine were analyzed in the urine collected before and after STZ injection using metabolic cages as described above.

### Renal morphology

At the end of experiments, mice were anesthetized and perfused with Dulbecco’s Phosphate buffered saline (PBS) pH 7.2 (Sigma, USA). The kidney was fixed in Formaldehyde 10% (v/v) buffered pH 7.4 Carson-Millonig (Rica, USA) for morphological analyses. Coronal sections of 3µ were stained with standard Hematoxylin-Eosin and Masson’s Trichrome and visualized by light microscopy. Also, kidney tissue was collected in CryoStor CS10 cell media (Sigma, USA) and cryopreserved in liquid nitrogen until further examination.

### Electron microscopy

Paraffin sections of 20µm thick kidney tissue were examined using transmission electron microscopy and the glomerular basement membrane thickness was measured as previously described.

### Data analysis

Data was analyzed using GraphPad Prism. For measured parameters, outliers were identified and removed using the Robust regression and outlier removal (ROUT) method with a false discovery rate of less than 1%. Based on this, four outliers were removed for albumin to creatinine ratio (ACR) that included one sample from the wild-type group at baseline and at endpoint each and two samples from *fcer1g*^*-/-*^ group at endpoint. Similarly, two data points were removed as outliers for glucose measurement in wild type at baseline.

## Results

### Both *fcer1^+/+^* and *fcer1^-/-^* mice are hyperglycemic in response to STZ

An earlier study identified FcER1 to be elevated in DKD and its expression was positively correlated with the disease state [7]. Therefore, we set out to test the efficacy of inhibition of FcER1 signaling in preventing or slowing the progression of DKD as outlined in the study design (**Figure 1**). First, we evaluated the effect of genetic deletion of *fcer1* on the development of DKD in response to STZ in a mouse model (*fcer1*^*-/-*^ mice). *Fcer1*^*-/-*^ mice were developed by Takai et al by deletion of the gamma chain subunit resulting in defects in NK cell mediated antibody dependent cytotoxicity and mast cell mediated allergic responses but no renal phenotype was reported [13].

Wild type and *fcer1*^*-/-*^ mice at 9 weeks of age were administered STZ to induce type I diabetes phenotype and associated renal injury as described in materials and methods (**Figure 2A**). The initial body weight was not different between the two groups, however the baseline glucose and albumin to creatinine (ACR) was slightly but significantly elevated in *fcer1*^*-/-*^ mice (**Figure S1A-D)**. The *fcer1*^*-/-*^ mice responded to STZ more rapidly with significantly higher blood glucose and urine excretion at 1 week which equalized over the course of the experiment (**Figure 2B and C**). Some mice presented with weight loss, apathy and hunchback earlier in the treatment and required euthanasia as per UCSF regulations. This was higher in *fcer1*^*-/-*^ mice (approximately 40% of knockout mice presented with symptoms in the first four weeks after STZ administration versus wild type mice where the presentation was much later) presumably due to the rapid elevation in blood glucose. Therefore, at the study end point (12-13 weeks after STZ administration) 13 wild type and 10 knockout mice were examined for their renal function in the context of hyperglycemia. There was no significant difference in food and water consumption during the experiment between the groups (**Figure S1E and 1F**). Blood glucose and urine volume were not different between wild type and *fcer1*^*-/-*^ mice at the end of experiment (**Figure 2B and 2C**).

**Figure 2.**
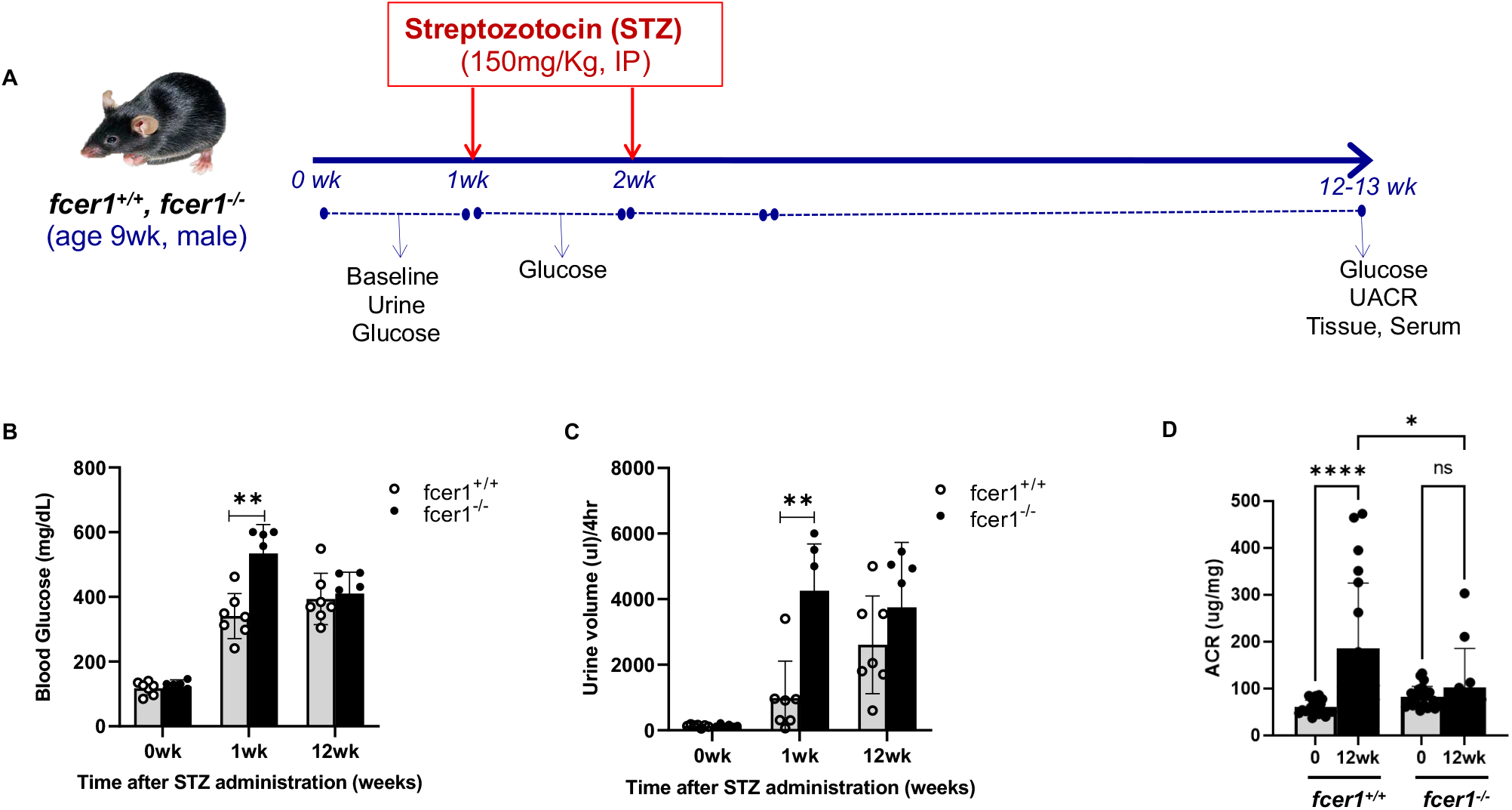
Streptozotocin (STZ) treatment of *fcer1*^*+/+*^ and *fcer1*^*-/-*^ mice. A) Schematic outlining the study design. B) Blood glucose before (baseline) and after 1 week and 12 weeks of STZ treatment. C) Urine excretion before (Baseline) and after 1 week and 12 weeks of STZ treatment. D) Albumin to creatinine ratio (ACR) is significantly lower in *fcer1*^*-/-*^ mice after STZ treatment compared to that in the *fcer1*^*+/+*^ mice.

### *fcer1^-/-^* mice are protected from STZ induced elevation in ACR

As described in methods, urine was collected from all mice before and after STZ administration. Urinary albumin and creatinine were measured on each sample on the same day and albumin to creatinine ratio (ACR) in the urine was calculated. STZ administration in *fcer1*^*+/+*^ mice caused approximately 2-3-fold increase in albuminuria 12-13 weeks after STZ injection while the knockout mice had no significant increase in urinary ACR (**Figure 2D**). At the end of 12 weeks, ACR was significantly higher in *fcer1*^*+/+*^ mice compared with the *fcer1*^*-/-*^ mice. Two-way ANOVA analysis showed that the absence of *fcer1* had a significant effect on ACR after STZ administration (p value <0.0001). At the end of the experiment, serum electrolytes and cholesterol levels were comparable between the STZ treated *fcer1*^*+/+*^ and *fcer1*^*-/-*^ mice except for serum sodium which was slightly but significantly lower in *fcer1*^*-/-*^ mice (**Table 1**). Serum albumin was also slightly but significantly lower in *fcer1*^*-/-*^ mice consistent with improved renal function compared to wild type mice after induction of diabetes (**Table 1**). At baseline, the glucose excretion in urine was slightly but significantly reduced and phosphorous excretion was elevated at the end of study in *fcer1*^*-/-*^ mice (**Table 2**).

**Table 1.** Serum parameters of *fcer1*^*+/+*^ and *fcer1*^*-/-*^ mice 12-13 weeks after STZ injection (study end). Serum parameters for *fcer1*^*+/+*^ and *fcer1*^*-/-*^ mice were compared using Students’ t-=test. ** p value<0.05. HDL: High-density lipoprotein; LDL: Low-density lipoprotein; BUN: Blood urea nitrogen; Cr: Creatinine; Ca: Calcium; Phos: phosphorous; Na: Sodium; K: Potassium; Cl: Chloride.

| Serum parameters | *fcer1<sup>+/+</sup>* | *fcer1<sup>-/-</sup>* | p value |
| --- | --- | --- | --- |
| Glucose (mg/dL) | 592.75±88.1 | 961.5±161.93 | 0.15 |
| Cholesterol (mg/dL) | 97.25±3.5 | 114.5±20.04 | 0.18 |
| Triglycerides (mg/dL) | 126.75±3.5 | 184.25±120.51 | 0.41 |
| HDL (mg/dL) | 83±3.6 | 99±15.1 | 0.12 |
| LDL (mg/dL) | 5±0.82 | 5±1 | 1.00 |
| BUN (mg/dL) | 37.5±4.8 | 45.5±9.19 | 0.42 |
| Cr (mg/dL) | 0.42±0.04 | 0.4575±0.24 | 0.79 |
| Ca (mg/dL) | 6.83±1.8 | 8.8±0.28 | 0.12 |
| Phos (mg/dL) | 7.1±0.76 | 6.25±0.07 | 0.11 |
| Total protein (g/dL) | 5.1±0.2 | 5.0±0.2 | 0.49 |
| Albumin (g/dL) | 2.9±0.1 | 2.6±0 | **0.003\*\*** |
| Na (mmol/L) | 158.5±3.32 | 150.5±0.71 | **0.013\*\*** |
| K (mmol/L) | 4.6±0.29 | 4.75±0.35 | 0.66 |
| Cl (mmol/L) | 108.5±6.14 | 102±0 | 0.12 |
| CO2 (mmol/L) | 18.6±1.77 | 18.4±0.99 | 0.87 |
| Na/K ratio | 34.58±2.53 | 31.8±2.55 | 0.33 |
| Anion gap (mmol/L) | 36±8.23 | 34.85±0.64 | 0.80 |

**Table 2.** Urinary parameters of fcer1g+/+ and fcer1g-/-mice at the start of the experiment and 12-13 weeks after induction of diabetes with STZ injection (study end). Parameters for Fcer1g+/+ and Fcer1g-/-mice were compared using Students’ t-test. ** p value<0.05. UUN: Urinary urea nitrogen; Phos: phosphorous; Ca: Calcium; Na: Sodium; K: Potassium; Cl: Chloride; Mg: Magnesium.

| Urinary parameters | Baseline |  |  | +STZ |  |  |
| --- | --- | --- | --- | --- | --- | --- |
|  | *fcer1*<sup>+/+</sup> | *fcer1*<sup>-/-</sup> | p value | *fcer1*<sup>+/+</sup> | *fcer1*<sup>-/-</sup> | p value |
| Glucose (mg/dL) | 177±37.63 | 140.71±23.21 | **0.04** | 8687.6±2571.97 | 9178±1072.28 | 0.518 |
| BUN (mg/dL) | 5711.375±524.06 | 5235.57±909.25 | 0.25 | 756.57±212.66 | 650.9±108.15 | 0.127 |
| Phosphorous (mg/dL) | 253.87±20.86 | 249.34±57.03 | 0.85 | 11.28±8.71 | 20.64±10.61 | **0.045** |
| Ca (mg/dL) | 13.04±4.03 | 12.49±1.19 | 0.72 | 5.08±2.38 | 6.08±3.54 | 0.475 |
| Na (mmol/L) | 196.33±92.35 | 141.71±11.7 | 0.14 | 50.11±12.91 | 40.16±20.46 | 0.196 |
| K (mmol/L) | 581.98±111.3 | 466.79±70.14 | **0.03** | 67.31±15.04 | 58.22±10.27 | 0.092 |
| Cl (mmol/L) | 467.6±117.43 | 368.83±66.47 | 0.07 | 65.54±17.29 | 60.11±18.01 | 0.468 |
| Mg (mmol/L) | 96.55±13.11 | 86.84±14.55 | 0.20 | 14.78±4.17 | 11.41±4.26 | 0.069 |

### Histopathology and morphometric analysis show attenuation of alterations in renal tissue of *fcer1^-/-^* mice

To assess the renal injury in STZ treated mice, tissue sections were examined by light microscopy after Gomori’s trichrome **(Figure 3A**) and hemotoxin and eosin (H&E) (**Figure 3C**) staining. Kidneys from *fcer1*^*+/+*^ mice show significantly higher perivascular fibrosis (**Figure 3B**), and perivascular inflammatory cells infiltrate (**Figure 3A-C**). Also, cortical interstitial fibrosis and cortical lymphocytic infiltrate were observed preferentially in kidney tissue of *fcer1*^*-/-*^ mice. Total number of inflammatory cells clusters (**Figure 3D**) as well as specifically mast cells as visualized by staining with a monoclonal antibody to tryptase (**Figure 3E**) was significantly lower in the *fcer1*^*-/-*^ kidney (**Figure 3F**). Both *fcer1*^*+/+*^ and *fcer1*^*-/-*^ mice showed discrete ultrastructural changes after STZ administration such as mild irregular thickening of the glomerular basement membrane. There were also rare bridges between the capillary tuft and Bowman’s capsule (**Figure S2**). Mild coarsening and effacement of podocyte foot processes were observed in *fcer1*^*+/+*^ mice, while *fcer1*^*-/-*^ mice maintained normal podocyte architecture after STZ treatment (**Figure 3G**). The alterations observed are described in very early stages of diabetic kidney disease [14, 15].

**Figure 3.**
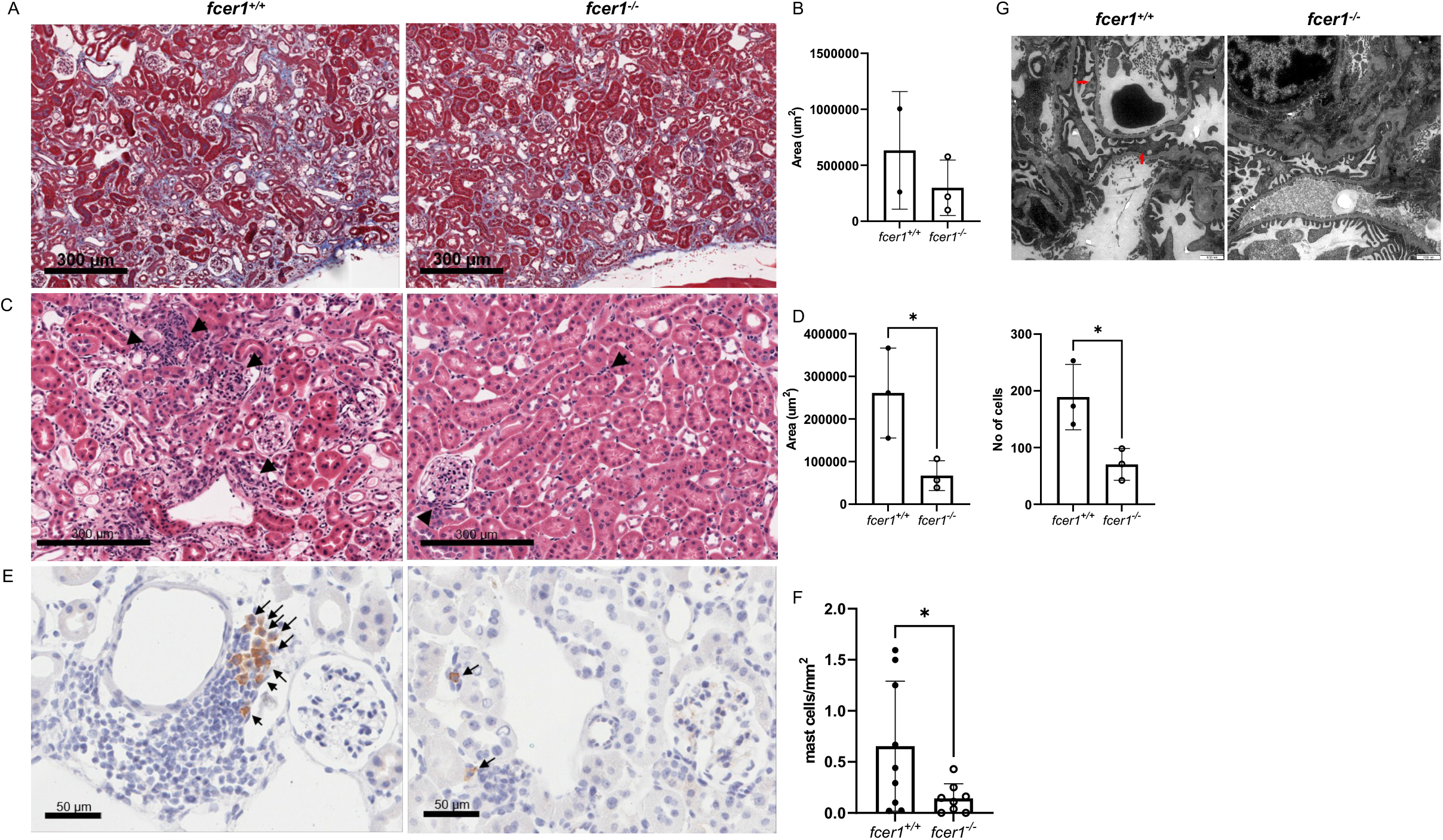
A) Representative image of Trichome staining of renal tissue from *fcer1*^*+/+*^ and *fcer1*^*-/-*^ mice shows fibrosis. B) Quantification of fibrosis shows significantly reduced fibrosis in *fcer1*^*-/-*^ mice compared to that in the *fcer1*^*+/+*^. C) Hematoxin and Eosin (H&E) staining of kidney from *fcer1*^*+/+*^ and *fcer1*^*-/-*^ mice. D) The areas of inflammation as well as the number of infiltrating cells is significantly reduced in *fcer1*^*-/-*^ mice after STZ treatment. E) A representative image of mast cell infiltration near the microvessels visualized by Tryptase staining. Arrows indicate tryptase^+^ mast cells. F) Quantification of mast cell number per mm^2^ in *fcer1*^*+/+*^ and *fcer1*^*-/-*^ mice shows significantly lower in *fcer1*^*-/-*^ mice. G) Podocyte foot process effacement in *fcer1*^*+/+*^ mice while *fcer1*^*-/-*^ mice retain their podocyte architecture.

### Inhibition of FcER1 receptor with an anti γ-subunit antibody does not prevent development of renal injury in response to STZ

Stabilization of mast cells has been shown to improve outcomes in obesity and diabetes [11]. In order to test the hypothesis that inhibition of the FcER1 receptor signaling before administration of STZ will prevent development of proteinuria in mice, we pre-treated C57BL6 wild type mice with two weekly doses of MAR-1 followed by one high dose STZ (Study design, **Figure 4A**). STZ induced hyperglymia and increased ACR in mice pre-treated with IgG control and MAR-1. At the experimental end-point, no difference in blood glucose or ACR were found between control IgG and MAR-1 treated mice (**Figure 4B and 4C**). We also did not see any structural alterations by light microscopy (**Figure 4D**).

**Figure 4.**
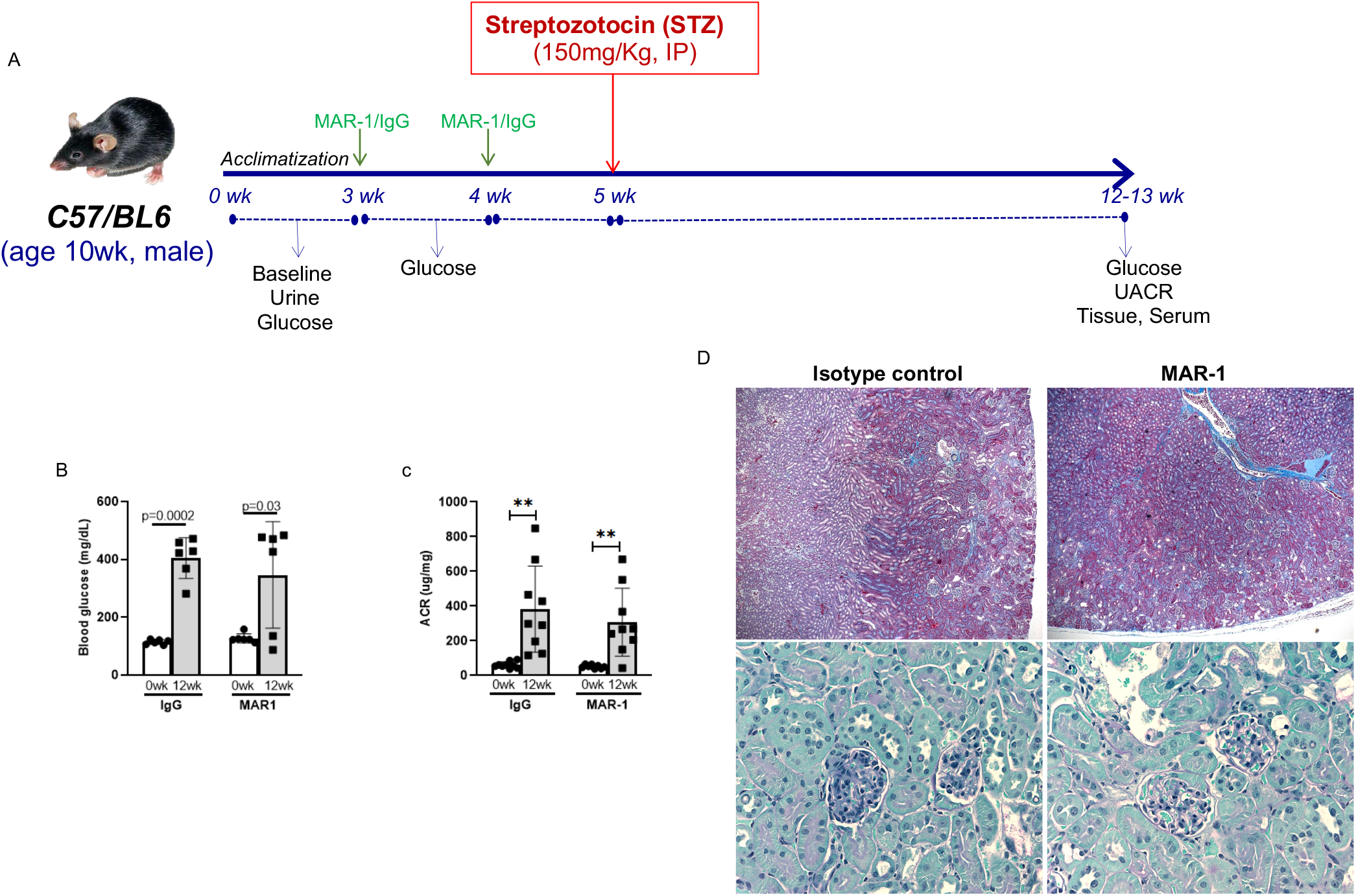
Pre-treatment of wild type C57BL6 mice with anti FcER1 antibody does not prevent the development of albuminuria after STZ treatment. A) Wild type C57BL6 mice were treated with two weekly doses of 50ug MAR-1 antibody (IP) followed by one high dose STZ injection to induce hyperglycemia and associated renal injury. Urine and blood were collected at the beginning and end of the study. Blood glucose levels were measured at the beginning and end of the study. B) Serum glucose (mg/dL) in mice treated with IgG control or MAR-1 antibody at the beginning (0wk) and end (12wk) of study. C) Albumin to Creatinine ration in urine of mice treated with isotype IgG control (ug/mg). D) Histological analysis of wild type C57BL6 mice treated with IgG isotype control or MAR-1 antibody before STZ administration to induce hyperglycemia and proteinuria shows no difference in fibrosis (top, Gomori trichrome staining) and inflammatory infiltrates (bottom, PAS staining).

### Inhibition of FcER1 receptor with an anti γ-subunit antibody attenuates the development of proteinuria in a spontaneous diabetes mouse model

To evaluate the therapeutic benefit of FcER1 antagonism in diabetic kidney disease in a mouse model that spontaneously develops diabetes, we chose BKS.Cg-+Leprdb/+Leprdb/OlaHsd (*db/db*) male mice (The Jackson Lab) homozygous for the diabetes spontaneous mutation in the leptin receptor gene (*Lepr*^*db*^). At 16weeks old, *db/db* mice have hyperglycemia and diabetes associated renal injury is established. Mice were treated weekly with 50ug MAR-1 or an isotype control for 6 weeks as described in materials and methods. As control, one group of mice were left untreated (see study design, **Figure 5A**). No significant differences were observed between mice injected with MAR-1 and untreated mice (data not shown). We did not see a significant difference between body weights of untreated, IgG treated or MAR-1 treated mice during the course of the study (**Figure S3**). As previously described, mice with blood glucose levels greater than 250 mg/dl were confirmed as diabetic [16]. All mice were diabetic at the beginning and end of study. There is a small but significant increase in blood glucose level in MAR-1 treated mice at the end of treatment (**Figure 5B**). However, there were no significant differences in the blood glucose level between MAR-1 and IgG treated mice at any time point. 24-Hour Urine samples of all animals were collected at the beginning and the end of the study using rodent metabolic chambers as described in materials and methods. The urine volume remained unchanged between control and various treatment groups (data not shown). On the other hand, albuminuria as represented by urinary albumin normalized to creatinine (ACR) significantly increased in IgG control group of mice while the MAR1 treated mice remain protected (**Figure 5C**). At the end of 6week after MAR1 treatment, mice had significantly lower ACR than that of the IgG control mice (**Figure 5C**). Histological analysis of renal tissue from *db/db* mice at the end of treatment showed moderate fibrosis in both cortical and medullary areas of the kidney along with few inflammatory infiltrates, however, no significant differences in fibrosis or inflammation between IgG and MAR-1 treated groups were observed (**Figure 5D**). We did not observe any significant differences in serum electrolytes, BUN or serum creatinine between IgG and MAR-1 treatments. However, MAR-1 administration resulted in significant reduction in the excretion of phosphorous as well as K, Cl and Mg electrolytes (**Table 3**).

**Figure 5.**
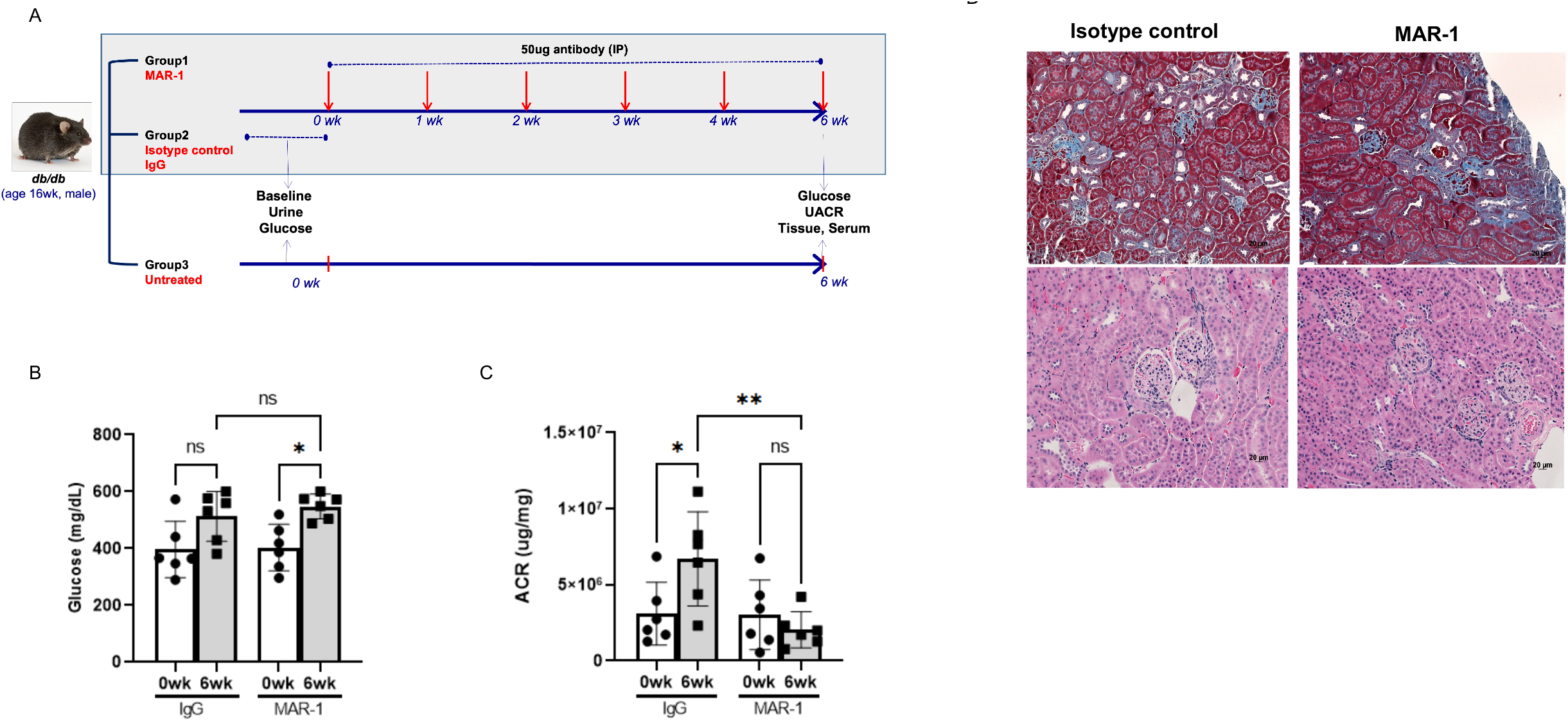
Inhibition of FcER1 signaling with anti FcER1 antibody attenuates the progression of albuminuria in a *db/db* mouse model of spontaneous diabetes. A) 16 week old mice were treated with MAR-1 antibody (Group1, n=6), isotype control IgG (Group 2, n=6) or left untreated (Group3, n=5). 50ug of antibody was weekly injected intraperitoneally (IP) for six weeks. At the end of the experiment, urine, serum and renal tissue was collected. Serum glucose was measured at baseline and at the end of the experiment. B) Serum glucose (mg/dL) in *db/db* mice treated with IgG control or MAR-1 at the start and end of experiment. C) Urinary albumin to creatinine (ACR, ug/mg) shows significant increase in proteinuria in control mice but not in MAR-1 treated mice. D) A representative image of db/db mouse kidneys after treatment with isotype control or MAR-1 and stained with Gomori’s trichrome to visualize fibrosis (top) and Hematoxin and Eosin (H&E) to visualize inflammatory infiltrates (bottom). The images are representative of 3-5 mice in each group.

**Table 3.**
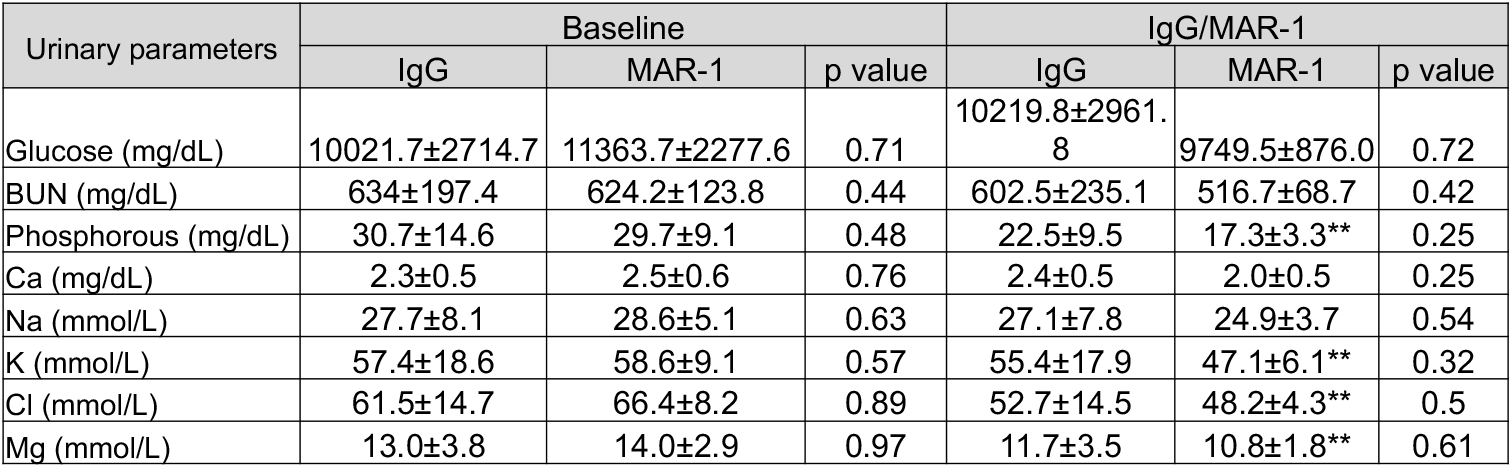
Urinary parameters of *db/db* mice at the start of the experiment and after MAR-1 injection (study end). Parameters for mice injected with IgG or MAR-1 were compared using Students’ t-test and p values for each parameter between IgG and MAR-1 injected mice at baseline or after IgG/MAR-1 injection are listed. ** p value<0.05 with respect to baseline for the treatment indicated. UUN: Urinary urea nitrogen; Phos: phosphorous; Ca: Calcium; Na: Sodium; K: Potassium; Cl: Chloride; Mg: Magnesium.

| Urinary parameters | Baseline |  |  | IgG/MAR-1 |  |  |
| --- | --- | --- | --- | --- | --- | --- |
|  | IgG | MAR-1 | p value | IgG | MAR-1 | p value |
| Glucose (mg/dL) | 10021.7±2714.7 | 11363.7±2277.6 | 0.71 | 10219.8±2961.8 | 9749.5±876.0 | 0.72 |
| BUN (mg/dL) | 634±197.4 | 624.2±123.8 | 0.44 | 602.5±235.1 | 516.7±68.7 | 0.42 |
| Phosphorous (mg/dL) | 30.7±14.6 | 29.7±9.1 | 0.48 | 22.5±9.5 | 17.3±3.3** | 0.25 |
| Ca (mg/dL) | 2.3±0.5 | 2.5±0.6 | 0.76 | 2.4±0.5 | 2.0±0.5 | 0.25 |
| Na (mmol/L) | 27.7±8.1 | 28.6±5.1 | 0.63 | 27.1±7.8 | 24.9±3.7 | 0.54 |
| K (mmol/L) | 57.4±18.6 | 58.6±9.1 | 0.57 | 55.4±17.9 | 47.1±6.1** | 0.32 |
| Cl (mmol/L) | 61.5±14.7 | 66.4±8.2 | 0.89 | 52.7±14.5 | 48.2±4.3** | 0.5 |
| Mg (mmol/L) | 13.0±3.8 | 14.0±2.9 | 0.97 | 11.7±3.5 | 10.8±1.8** | 0.61 |

## Discussion

Diabetic kidney disease is the most common cause of end stage renal disease. While diet and therapeutic control of glycemia and blood pressure has resulted in significant reduction in cardiovascular complications associated with diabetes, little effect has been observed on renal injury. Currently there are no direct therapeutic drugs available to treat or slow down the progression of renal injury in diabetes. In this article we show for the first time that *fcer1* deletion in mice protects them from renal injury in a mouse model of streptozotocin induced hyperglycemia. In this model, induction of diabetes with STZ did not lead to changes in the initial body weight; however, ACR was significantly lower in *fcer1*^*-/-*^ mice. Using STZ-induced diabetes in the *fcer1*^*-/-*^ model we observe a slight but significant increase in sensitivity to STZ in *fcer1*^-/-^ mice that could be due to higher susceptibility of pancreatic beta cells to STZ or altered insulin production mice. Approximately 40% of knockout mice presented with symptoms in the first four weeks after STZ administration versus wild type mice where the presentation was much later) presumably due to the rapid elevation in blood glucose. The altered sensitivity to glucose could in fact be due to mast cells which have been implicated in the development of severe insulitis and accelerated hyperglycemia [17]. Individuals with type I or II diabetes have accumulation of mast cells in the pancreas [18]. While DKD is not a primary inflammatory disease, there is large evidence of participation of inflammatory cells in DKD [19]. In in *vivo* models of DKD the Inflammation has been shown to be involved in the pathogenesis of CKD and is accompanied with the immediate release of histamine and synthesis of prostaglandins and leukotrienes by activated mast cells expressing FcER1. Subsequently, it stimulates the delayed secretion of cytokine and chemokine contributing to the inflammatory milieu around the renal lesion in diabetes [20]. Mast cells are found in tubulointerstitial and glomerular regions of kidney in association with T cells [19]. In order to understand the relationship between FcER1 mediated inflammation and renal function, we studied the renal morphology and presence of inflammatory molecules in STZ treated mice. The animals showed drastic reduction of mast cells (as determined by positive tryptase staining) in the *fcer1*^*-/-*^ kidney and were protected from tubular and glomerular injury as well as proteinuria development. Mast cells were observed in cortical and glomerular regions of kidney after STZ treatment. Interestingly histopathology analysis showed that fcer*1*^*-/-*^ mice had reduced number of inflammatory infiltrates in medullary and cortical kidney tissue after STZ treatment further supporting the role of inflammatory mediators in DKD pathogenesis. The pathway involving mast cells and inflammation in the progression of DKD is complex but it is proposed that the activation of mast cell could favor the recruitment of inflammatory cells into kidney lesions. In fact, anti-inflammatory therapies as such inhibition of NF-κB, AMPK, TLRs, MyD88 or HIF signaling pathways, inflammasome activation, mtDNA release can help in the reduction of inflammation-induced renal damage of DKD [21]. At this time, very few drugs have been approved by the FDA to treat nephropathy in diabetic individuals. We hypothesized that early treatment of pre-diabetic patients with antibodies recognizing FcER1 could inhibit the progression of DKD without altering their glycemic status. We treated wild type C57BL6 mice with two doses of MAR-1 to inhibit FcER1 signaling before STZ mediated induction of hyperglycemia. However, we did not observe an effect on proteinuria or renal morphology. Further studies with altered duration (such as MAR-1 treatment throughout the experiment) or dose of MAR-1 would be required to conclusively test the hypothesis.

To explore the immunotherapeutic potential of FcER1 inhibition with MAR-1 antibody concomitant with progression of diabetes we weekly administered MAR-1 in a mouse model of type II diabetes with established hyperglycemia and proteinuria. At 21 weeks of treatment, proteinuria was significantly lower in MAR-1 treated mice but we observed no evidence of improvement in fibrosis, inflammation or in the histopathologic score. It is possible that a longer inhibition of FcER1 signaling is required to achieve reversal of histological injury, however a prolonged treatment with antibody poses a risk of immunological response against the antibody. In such scenario, humanized antibody will be a good alternative [22]. At present, there is no antibody available in the clinic to perturb FcER1 signaling, therefore, we conducted the study with MAR-1 which is developed by eBiosciences (now part of Thermo Scientific) and reacts with the Fc epsilon receptor I alpha subunit. This subunit binds to IgE but lacks any signal transducing ability. Therefore, pharmacological disruption of FcER1 signaling by MAR-1 is currently not well understood. Several studies have demonstrated that in addition to inhibiting FcER1, MAR-1 also cross-reacts with FcγRI and FcγRIV [23]. The immunomodulation using anti-Fc or anti-receptor antibodies with high specificity to only FcER1 is more complex due to cross-linking between other Fc receptors on mast cells and basophil. Future studies focusing on new strategies of designing new antibodies and pharmacological approaches to confirm the ability of FcER1 in modulating the progression of DKD are needed.

In conclusion, we have shown that FcER1 receptor plays an essential role in the development of renal injury associated with diabetes and the inhibition of FcER1 signaling mediated inflammation in kidney has therapeutic potential to treat DKD.

## Figure Legends

**Figure S1**. A) Body weight of *fcer1*^*+/+*^ (n=26) and *fcer1*^*-/-*^ mice (n=35) was not different at the start of the experiment. B) Baseline glucose was observed to be slightly but significantly higher in *fcer1*^*-/-*^ mice (n=36) versus *fcer1*^*+/+*^ (n=32). C) Baseline urine volume was not different in *fcer1*^*-/-*^ mice (n=17) from *fcer1*^*+/+*^ (n=20) mice. D) Baseline ACR (ug/mg) was significantly higher in *fcer1*^*-/-*^ mice compared to *fcer1*^*+/+*^ control. E) Body weight (g), F) Blood glucose (mg/dL), and G) Urine volume (μl) during the experiment in *fcer1*^*+/+*^ and *fcer1*^*-/-*^ mice. Data was analyzed using non-parametric Mann-Whitney test.

**Figure S2**. Electron microscopy shows formation of bridges in *fcer1*^*+/+*^ mice that are absent in *fcer1*^*-/-*^ *mice*.

**Figure S3**. A) Body weight of *db/db* mice were measured weekly (g) through the course of experiment before administration of IgG or MAR-1. B) Urine volume (mL) at the beginning and end of experiment.

## Acknowledgements

This study was supported by National Institute of Health grant R21 TR001761 01.

## Notes

### Competing Interest Statement

The authors have declared no competing interest.

