## Supplementary figures and images for "Genetic deletion and pharmacologic inhibition of FcER1 reduces renal injury in mouse models of diabetic nephropathy"

### Supplemental Figures S1-S3

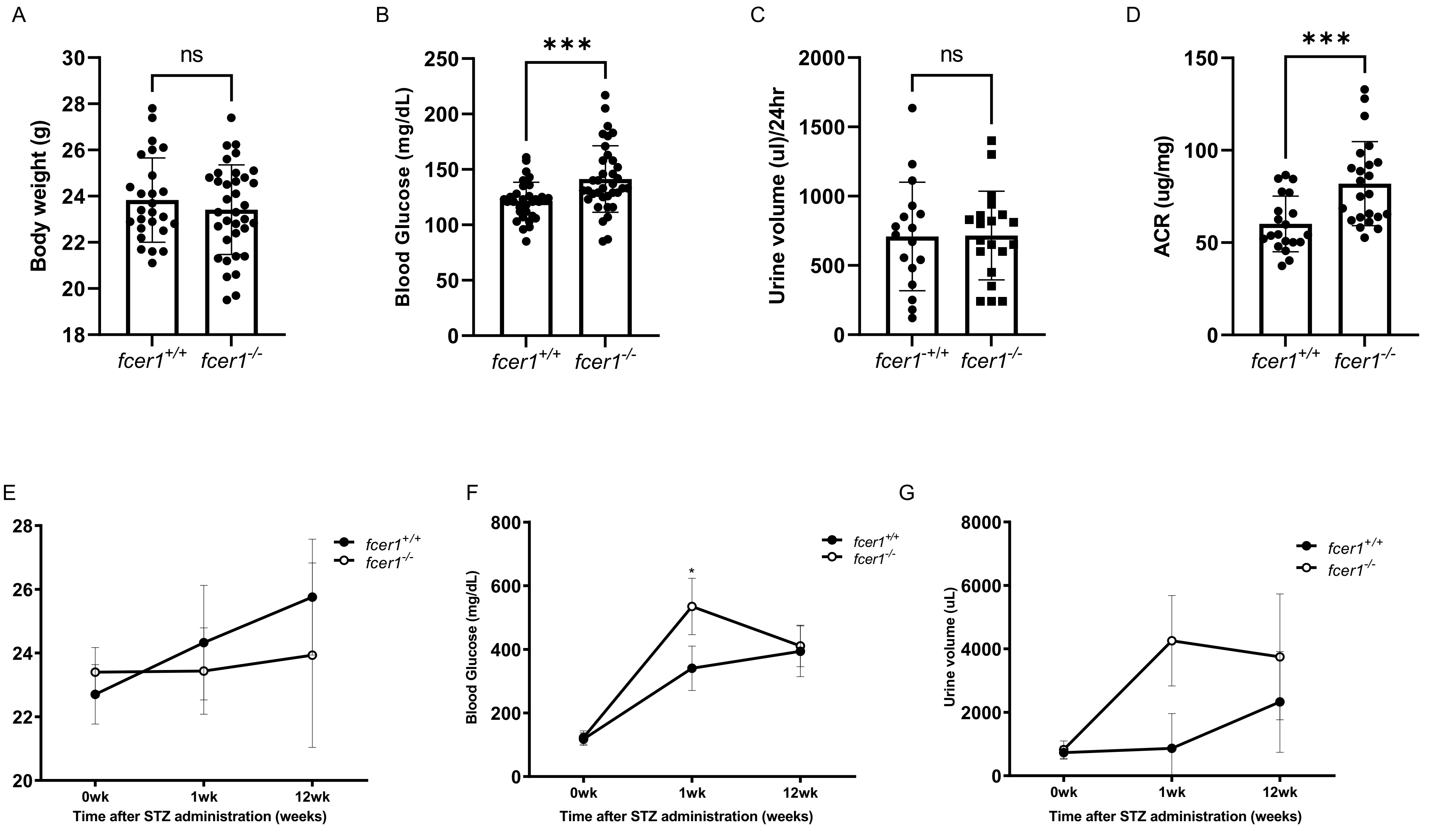

Figure S2

***fcer1***$^{+/+}$ ***fcer1***$^{-/-}$

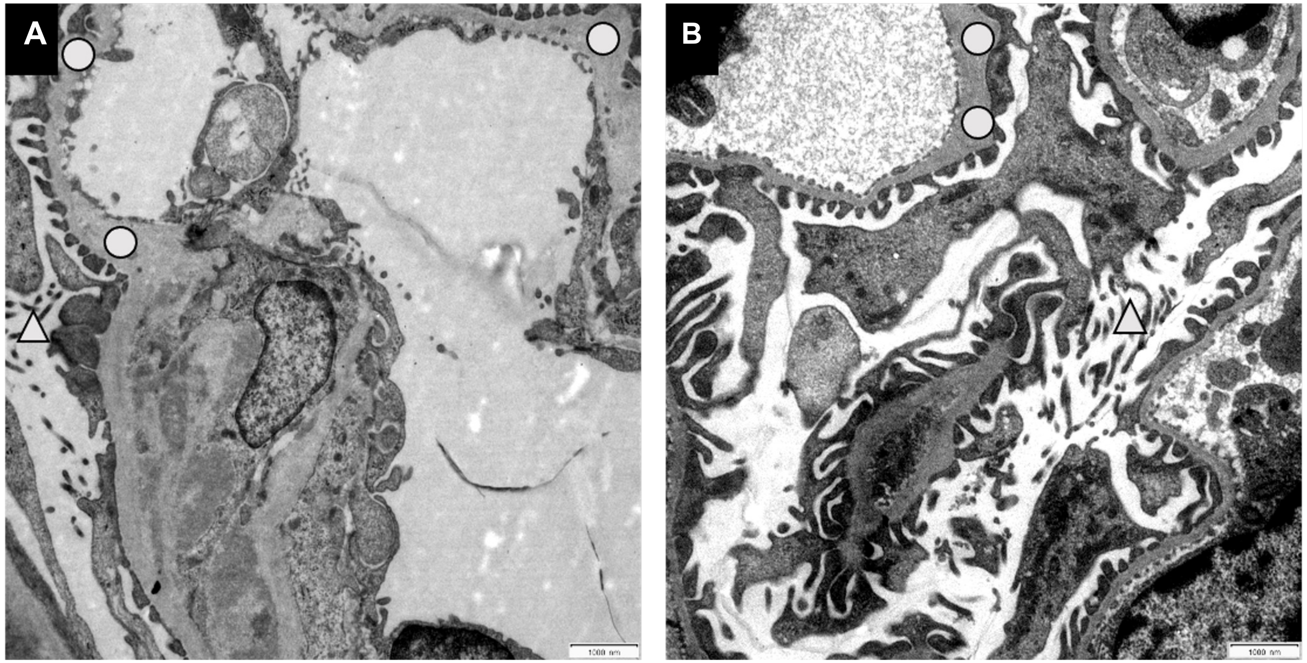

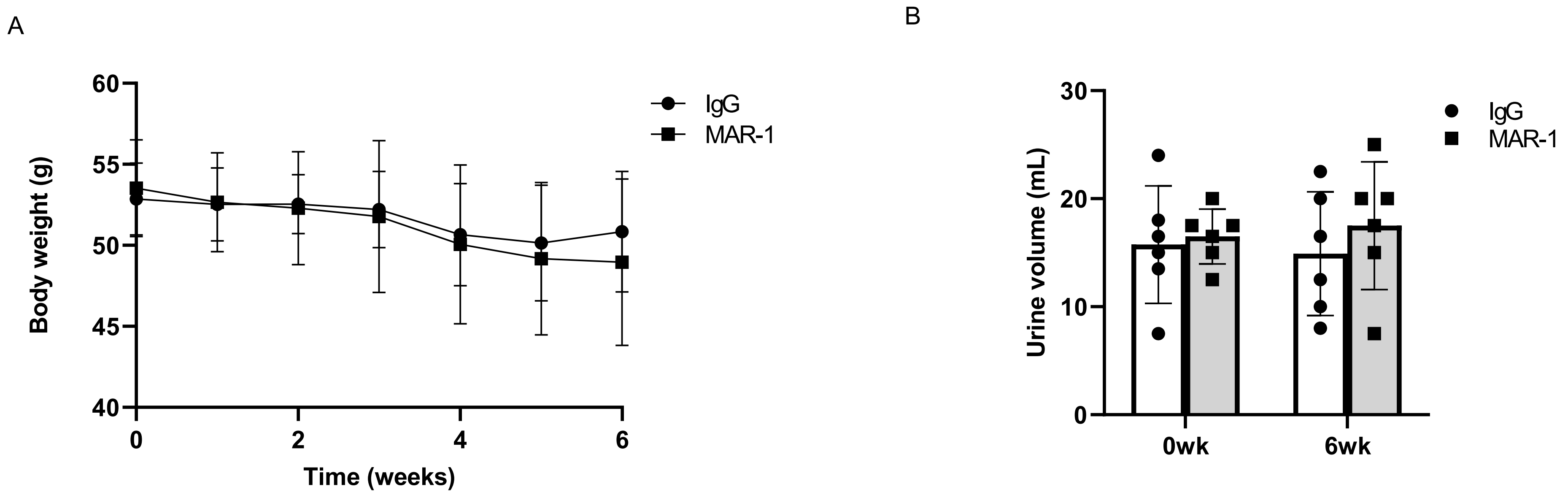
